# Thesaurus: quantifying phosphoprotein positional isomers

**DOI:** 10.1101/421214

**Authors:** Brian C. Searle, Robert T. Lawrence, Michael J. MacCoss, Judit Villén

## Abstract

Proteins can be phosphorylated at neighboring sites resulting in different functional states, and studying the regulation of these sites has been challenging. Here we present Thesaurus, a search engine that detects new positional isomers using site-specific fragment ions from parallel reaction monitoring and data independent acquisition mass spectrometry experiments. We apply Thesaurus to analyze phosphorylation events in the PI3K/AKT signaling pathway and show neighboring sites with distinct quantitative profiles, indicating regulation by different kinases.

---

It is estimated that hundreds of thousands of residues from thousands of proteins are actively phosphorylated in every human cell (1). Many proteins are phosphorylated at neighboring sites (2) where over half of sites in multi-phosphorylated proteins are within four amino acids of each other (3). Several well-studied proteins make use of adjacent phosphorylation sites to act as switches (MAPK (4), CDC4 (5)), timers (PER (6)) or as negative inhibition toggles (IRS1 (7)) but global analysis of these phosphorylation clusters has remained impractical. Tandem mass spectrometry (MS/MS) of tryptic peptides is a key tool in discovering and quantifying of sites of protein phosphorylation. Typical phosphoproteomic workflows use data dependent acquisition (DDA) to collect MS/MS spectra based on peptide precursor m/z as peptides chromatographically elute. To increase the number of unique peptides that are sampled, DDA dynamically excludes peptides of the same m/z from being sampled twice within a narrow elution time. However, peptides that exist as multiple positional phosphoisomers have 1) the same mass, 2) similar retention times, and 3) many of the same fragment ions, making them extremely difficult to differentiate. Thus MS/MS sampling using DDA and dynamic exclusion makes it challenging to sample multiple positional isomers.

Parallel reaction monitoring (PRM) and data independent acquisition (DIA) represent an alternative class of acquisition approaches that systematically collect MS/MS spectra across the chromatographic elution of the peptide, improving quantitative reproducibility. While PRM methods target specific peptide precursors, DIA methods iteratively sweep across m/z windows to acquire MS/MS spectra irrespective of peptides m/z that have been sampled previously (8). These methods are free of both: intensity biases during data collection and active exclusion of previously sequenced m/z, making it possible to detect closely eluting positional isomers. In addition, quantification can be performed on the more sensitive and selective product ion chromatograms, including those that are specific to each isomer.

Despite the strengths of these methods, mapping a phosphorylation site to a specific residue remains difficult. Rosenberger et al (9) proposed a method for determining the predominant modification site after library-search detection that determines the most likely peptidoform from fragment ions in a peak. An alternate approach (10) is to deconvolute DIA data (11) for processing with site localizing tools originally designed for DDA (12,13). Phosphopeptide isomers frequently co-elute; and a limitation of both of these approaches is that multiple forms with similar retention times are competed against each other, where at most one will be detected. Here we extend these approaches and present a new DIA and PRM hybrid (i.e. spectrum library/database) search engine named Thesaurus, which is designed to specifically look for and quantify all detectable positional isomers, including those that are not found in the library and isomers that co-elute.

Thesaurus detects phosphopeptides in spectrum libraries and uses those detections as retention time anchors to iteratively find new positional isomers that share many of the same fragment ions but differ in their site-specific ions (Figure 1a). Thesaurus can detect multiple positional isomers at the same time point because it calculates localization probabilities directly using an interference distribution rather than by competing isomers against each other. Also unlike previous methods (9,10), if Thesaurus can confidently assign the outermost positional forms of peptides with >2 acceptor residues, it then iteratively searches for other forms with phosphosites more internal to the peptide using the shared ions as long as they localize to distinct retention times. To accomplish this, Thesaurus extracts site-specific fragment ion signals for each combinatorially potential phosphoisomer and calculates the probability that those ions would be observed by chance. Each ion has a unique likelihood of being interfered with across the experiment, and this probability of interference is highest with low M/Z ions (Supplementary Figure 1). Thesaurus calculates this background frequency distribution for each precursor isolation window since this probability is heavily skewed by peptide composition and the fragment ion mass tolerance. Peptides are scored as –log_10_(p) and localizations are considered significant if they pass a p-value threshold <0.01; Thesaurus further reevaluates these novel detections using FDR analysis with Percolator (14). If a combinatorial phosphoisomer is not present in our spectrum library, Thesaurus can generate a synthetic spectrum using an approach similar to SpectraST (15) by shifting fragment ions from known phosphoisomers to aid in the detection. One major advantage of analyzing phosphoproteomes with PRM and DIA is that each isoform Thesaurus finds can be quantified using site-specific ions even if the precursor signals are convolved. Thesaurus quantifies phosphopeptides by automatically determining peak boundaries based on the site-specific ions and choosing other fragment ions that fit the same shape.

**Figure 1.**
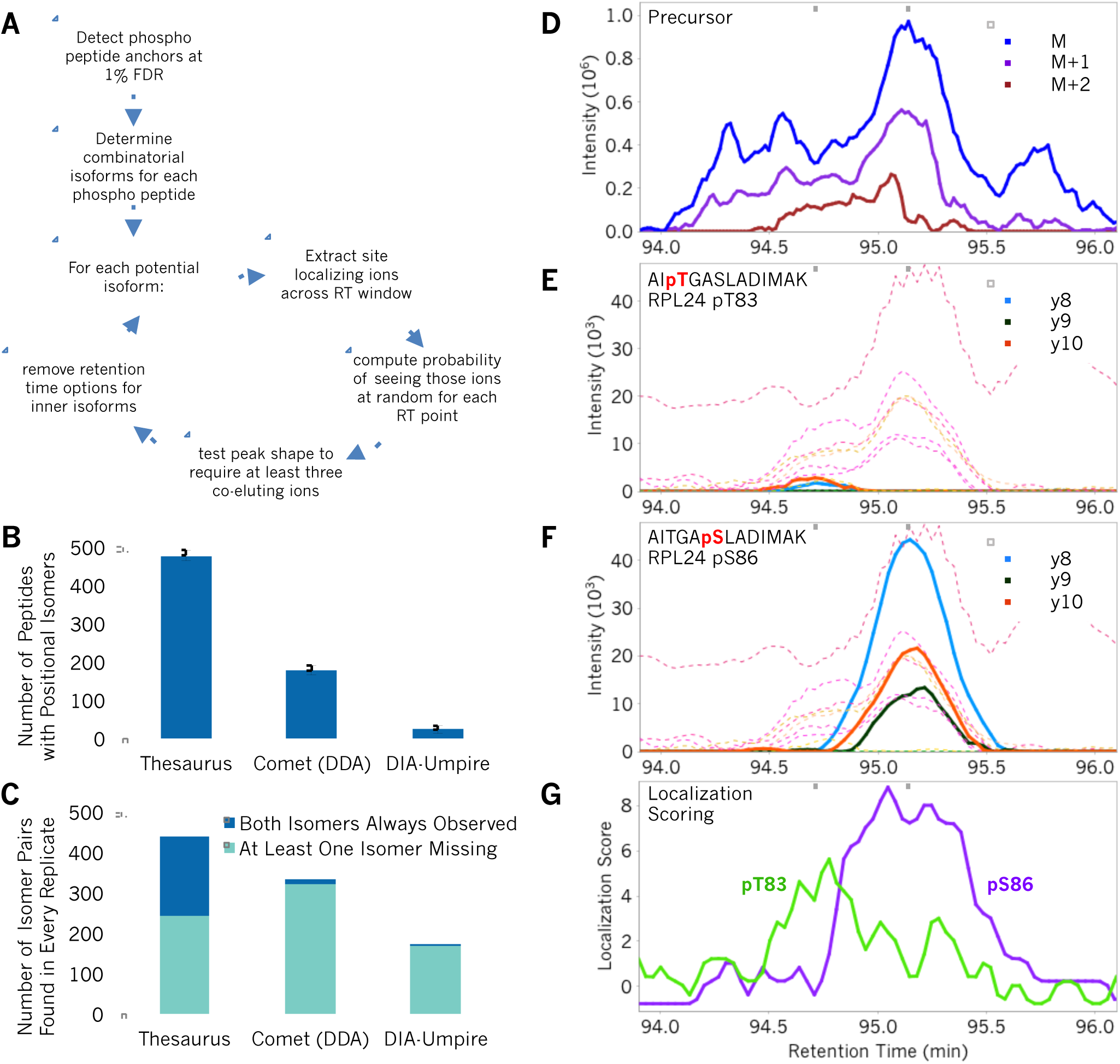
An approach for detecting phosphopeptides with Thesaurus. (**A**) Thesaurus algorithmic workflow to search for phosphoisomers from DIA data (see Online Methods). (B) The average number of peptides with multiple positional isomers that were detected from DIA data with Thesaurus and DIA-Umpire/Ascore, or DDA data with Comet/Ascore where error bars indicate 95% confidence intervals. (C) The number of singly phosphorylated peptides with two acceptor residues that were detected in all four technical replicates where both isomers were always observed or where an isomer was missing in least one replicate. To be included in this chart both forms of the peptide must have been observed in the same replicate by at least one analysis approach. (**D**) Precursor extracted ion chromatogram for the singly phosphorylated peptide AITGASLADIMAK from RPL24, which can be phosphorylated at both T83 and S86. Dashed grey lines indicate the retention time centers for the individual forms. Site-specific y8, y9, and y10 ions (solid) and other y-ions (dashed) for (**E**) pT83 and (**F**) pS86. (**G**) Localization scores for pT83 (green) and pS86 (purple).

To demonstrate the performance of our algorithm, we applied Thesaurus to phosphopeptides derived from serum-stimulated HeLa cells. Previously we reported a human phosphopeptide library based on nearly a thousand DDA experiments (16). In this work, we used a subset of this library that contains 82,029 phosphopeptides. Nearly half (44%) of these peptides could be localized to multiple forms (Supplementary Figure 2). Thesaurus was able to detect an average of 10,437 unique phosphopeptides across four technical replicate DIA experiments, corresponding to an average of 4,514 confidently localized isoforms (Supplementary Figure 3a). Approximately 22% of the peptides that contained multiple serines, threonines, or tyrosines could be localized to multiple positional isomers, and the retention time differences between these isomers were generally less than 60 seconds (Supplementary Figure 3b and 3c). We found that while Thesaurus performed comparably to Comet/Ascore with complementary DDA experiments (Supplementary Figure 4), Thesaurus found 2.7× more peptides with multiple positional isomers (Figure 1b) and 20× more than DIA-Umpire. In particular, when considering peptides with only two acceptor sites, Thesaurus was markedly better at consistently detecting both positional isomer pairs across all four replicates (Figure 1c). For example, in every replicate Thesaurus consistently detected two forms of the peptide AITGASLADIMAK from the 60S ribosomal protein RPL24 with single phosphorylation at T83 or S86. These forms co-elute within 25 seconds of each other, and while the precursor signal represents a mixture of both isomers (Figure 1d), Thesaurus confidently assigned them using site-specific ions (Figure 1e and 1f) to calculate localization scores (Figure 1g).

While DDA methods reliably triggered MS/MS on the less intense early eluting form (pS86), in 3 of the 4 DDA replicates MS/MS spectra corresponding to the more intense late eluting form (pT83) were never acquired due to dynamic exclusion. In these cases precursor quantification was unreliable because the signal from both forms was assigned exclusively to the low abundance form because it eluted first. This result suggests that run-to-run variability between phosphoproteome experiments might be due in part to a fundamental aspect of the data acquisition method. Using Comet (17) we were able to confirm that the site-specific fragment ions we observed in DIA are the same as those observed in the single DDA replicate that collected an MS/MS for this peptide (Supplementary Figure 5). In contrast, DIA-Umpire correctly generated a pseudo-MS2 for the higher intensity pT83 isoform in every DIA replicate but, due to interference, Comet scored the correct peptide only one of four times. In all four DIA replicates DIA-Umpire never generated a pseudo-MS2 for the lower intensity pS86 isoform.

Building on our new method to resolve phosphoisomers, we designed a DIA quantitative experiment to illuminate the PI3K/AKT signaling network in MCF-7 cells after stimulation with insulin and IGF-1. We found that 1,322 of the 5,944 localized and consistently measured phosphopeptides changed significantly at an FDR-corrected p-value <0.05 (Supplementary Table 1), where the predominant group showed increased abundance in insulin/IGF-1 regardless of MK-2206 treatment (Supplementary Figure 6a). As expected, some known AKT substrate sites showed significant response to MK-2206, such as FOXO3A S253 and PRAS40 T246 (Supplementary Figure 6b).

In this experiment, 28% of the localized phosphopeptides were observed as multiple positional isomers, and 23% of those showed >2-fold different induction from each other in response to insulin (Supplementary Figure 7). For example, Figure 2a diagrams how the peptide KGSGDYMPMSPK from the insulin receptor scaffold protein IRS1 contains three residues that are putatively phosphorylated by three different kinases: Y632 by INSR (upstream of AKT), S636 by S6K1 (downstream of AKT), and S629 by PKA. Our library had no spectral representation of KGSGDpYMPMSPK with which to make a spectrum library detection of phosphoY632. Despite this, Thesaurus was able to confidently detect, localize, and quantify all three singly phosphorylated forms (Figure 2b and 2c). We confirmed these detections with scheduled PRM runs on the same samples (Supplementary Figure 8) where the Thesaurus localization score can easily associate the retention times with each positional isomer. As expected from the model, after both insulin and IGF-1 stimulation phosphorylation of Y632 on INSR increased by >10-fold (Figure 2d). Similarly, S636 phosphorylation is also increased, but that effect was lower and significantly diminished when treated with the pan-AKT inhibitor MK-2206. As PKA is considered outside the AKT network, the phosphorylation state of S629 should stay unchanged and our measurements confirmed that. All three phosphopeptides contained several shared fragment ions and without statistical site localization it would have been extremely difficult and time consuming to manually determine which phosphoisomers were present and when they eluted. In addition to IRS1, we found numerous examples where only one phosphoisomer responded to insulin/IGF-1 stimulation or both responded with opposite directionality (Supplementary Figure 9). Finally, some phosphopeptide isomers were completely indistinguishable by retention time, yet could be confidently localized and quantified using site-specific ions. For example, MARK3 phosphoisomers at S469 and S476 co-eluted under our chromatographic conditions. Using Thesaurus, we were able to detect that the S469 isomer responded to insulin/IGF-1 and AKT inhibition while the S476 isomer remained constant (Supplementary Figure 10).

**Figure 2.**
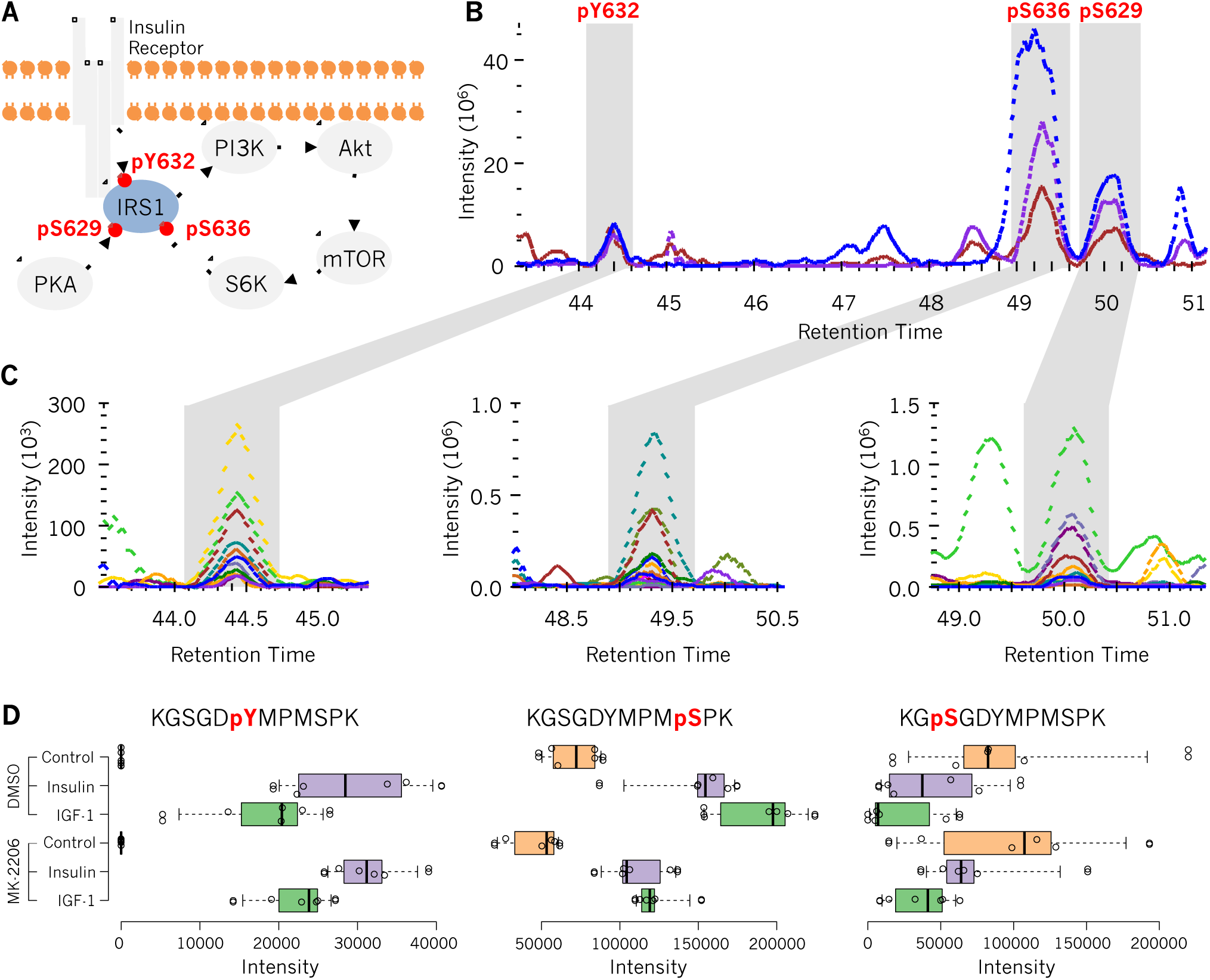
Detection and quantification of IRS1 phosphorylation. (**A**) Diagram of IRS1 phosphorylation at sites S629, Y632, and S636 with respect to insulin/IGF-1 stimulation and treatment with the AKT inhibitor MK-2206. (**B**) Precursor traces for three singly phosphorylated positional isomers of peptide KGSGDYMPMSPK from IRS1 in insulin-stimulated MCF-7 cells, and (**C**) corresponding fragment ions indicating phosphorylation at Y632 by INSR, S636 by S6K1, and S629 by PKA. Thesaurus was able to detect, localize, and quantify pY632 despite the peptide representing that form was absent from the searched spectrum library because it could use the pS629 and pS636 forms as anchors. (**D**) Box plots and values indicating summed fragment ion intensities for the three phosphosites on IRS1 across six replicates after stimulation with insulin, IGF-1, or unstimulated (control); with and without MK-2206. Boxes indicate quartiles and medians, while whiskers indicate the estimated 5% and 95% ranges.

Positional isomers represent an important concept for understanding signaling biology where neighboring phosphosites can have profound biological impact. Our tool Thesaurus provides a new avenue to study positional isoforms even if their precursor signals cannot be resolved. Now with a search engine specifically designed to analyze neighboring sites of phosphorylation it is possible to determine whether they have distinct functional implications, are redundant mechanisms for regulation, or are simply representative of a background phosphorylation state. The analysis of several other types of PTMs will also benefit from this analysis approach, so we have extended Thesaurus to support other modifications such as methylation, acetylation, and O-HexNAcylation and developed a robust, multithreaded tool with a stand-alone graphical interface to enable wide adoption. Our results indicate that PRM and DIA strategies will be crucial in assessing the complex regulatory nature of the human phosphoproteome.

## Methods

Methods are available in the online version of the paper.

## Acknowledgements

We would like to thank members of the Villén and MacCoss labs for critical discussions. B.C.S. is supported by F31 GM119273. This work is supported by grants P41 GM103533, R21 CA192983, and U54 HG008097 to M.J.M.; and R35 GM119536, R01 AG056359, and a research grant from the W.M. Keck Foundation to J.V.

## Author Contributions

R.T.L. and J.V. conceived the study. B.C.S., R.T.L., and J.V. designed the experiments. B.C.S. and R.T.L. performed the experiments. B.C.S. designed and wrote the software, and analyzed the data. M.J.M. and J.V. supervised the work. B.C.S., R.T.L., M.J.M. and J.V. wrote the paper.

## Online Methods

### Cell Culture

HeLa cervical cancer cells were cultured at 37°C and 5% CO_2_ in Dulbecco’s modified Eagle’s medium (DMEM) supplemented with L-glutamine, 10% FBS, and 0.5% streptomycin/penicillin. Cells were grown to an estimated 90% confluence in 10-cm plates, where one plate was used for each replicate/condition. Prior to harvest, cells were incubated for 4 hours under serum starvation conditions and then serum stimulated for 30 minutes. MCF-7 breast cancer cells were similarly cultured and starved, followed by stimulation with insulin (100 ng/ml) or IGF-1 (100 ng/ml) in phosphate-buffered saline (PBS) or unstimulated (control, added same volume of PBS) for 20 minutes. Some MCF-7 cells were additionally treated with DMSO or the pan-AKT inhibitor MK-2206 for 40 minutes before stimulation. After stimulation cells were quickly washed three times with refrigerated PBS and immediately flash frozen with liquid nitrogen. With the MCF-7 experiment, six replicates were performed for each of the six conditions: control/DMSO, insulin/DMSO, IGF-1/DMSO, control/MK-2206, insulin/MK-2206, and IGF-1/MK-2206. The six replicates were performed in three cell culture batches to simplify sample handling and ensure precise timing.

### Sample Preparation

Frozen cells were lysed in a buffer of 9 M urea, 50 mM Tris (pH 8), and 75 mM NaCl, with a cocktail of protease inhibitors (Roche Complete-mini EDTA-free) and phosphatase inhibitors (50 mM NaF, 50 mM β-glycerophosphate, 10 mM pyrophosphate, and 1 mM orthovanadate). After scraping, cells were subjected to 2 cycles of 25 seconds of probe sonication each followed by 10 minutes of incubation on ice. Lysates were centrifuged for 10 minutes at 21,000 × *g* and 4°C to eliminate cell debris. The protein content of the supernatant was estimated using BCA. For every condition/replicate, an estimated 850 μg of protein was reduced with 5 mM dithiothreitol for 30 minutes at 55°C, alkylated with 10 mM iodoacetamide in the dark for 30 minutes at room temperature, and the alkylation was quenched with an additional 5 mM dithiothreitol for 15 minutes at room temperature. The proteins were diluted to 1.8 M urea and then digested with sequencing grade trypsin (Pierce) at a 1:50 enzyme to substrate ratio for 4 hours at 37°C. The digestion was quenched by adding 10% trifluoroacetic acid (TFA) to achieve pH ~ 2. Resulting peptides were desalted with 100 mg tC18 SepPak cartridges (Waters) using vendor-provided protocols and dried with vacuum centrifugation. Phosphopeptides were enriched using immobilized metal affinity chromatography (IMAC) using Fe-NTA magnetic agarose beads (Cube Biotech). Enrichment was performed with a KingFisher Flex robot (Thermo Scientific), which incubated peptides with 150 μl 5% bead slurry in 80% acetonitrile, 0.1% TFA for 30 minutes, washed them three times with the same solution, and eluted them with 60 μl 50% acetonitrile: 1% NH_4_OH. Phosphopeptides were then acidified with 10% formic acid and dried. Phosphopeptides were brought to 1 μg / 3 μl in 0.1% formic acid assuming a 1:100 reduction in peptide abundance from the IMAC enrichment.

### Liquid Chromatography Mass Spectrometry

Phosphopeptides were separated with a Waters NanoAcquity UPLC and emitted into a Thermo Q-Exactive HF or a Thermo Fusion tandem mass spectrometer. Pulled tip columns were created from 75 um inner diameter fused silica capillary in-house using a laser pulling device and packed with 3 μm ReproSil-Pur C18 beads (Dr. Maisch) to 300 mm. Trap columns were created from 150 um inner diameter fused silica capillary fritted with Kasil on one end and packed with the same C18 beads to 25 mm. Solvent A was 0.1% formic acid in water, while solvent B was 0.1% formic acid in 98% acetonitrile. For each injection, 3 μl (approximately 1 μg) was loaded and eluted using a 90-minute gradient from 5 to 25% B, followed by a 40 minute washing gradient. Data were acquired using data-dependent acquisition (DDA), data-independent acquisition (DIA), or parallel reaction monitoring (PRM). Four DDA and DIA HeLa technical replicates were acquired in alternating mode to avoid bias. MCF-7 sample acquisition was randomized within blocks to enable downstream statistical analysis.

### DDA Acquisition and Processing

The Thermo Q-Exactive HF was set to positive mode in a top 12 configuration. Full MS scans of mass range 400-1600 were collected at 60,000 resolution to hit an AGC target of 3e6. The maximum inject time was set to 100 ms. MS/MS scans were collected at 30,000 resolution, AGC target of 1e6, and maximum inject time of 55 ms. The isolation width was set to 1.5 m/z with a normalized collision energy of 27. Only precursors charged between +2 and +4 that achieved a minimum AGC of 1e4 were acquired. Dynamic exclusion was set to “auto” and to exclude all isotopes in a cluster.

Thermo .RAW files were converted to .mzXML format using ReAdW and searched against a Uniprot Human FASTA database (downloaded July 1 2014 to maintain consistency with Lawrence *et al* (2), 87,613 entries) with Comet (version 2015.02v2), allowing for variable methionine oxidation, protein N-terminal acetylation, and phosphorylation at serines, threonines, and tyrosines. Cysteines were assumed to be fully carbamidomethylated. Searches were performed using a 50 ppm precursor tolerance and a 0.02 Da fragment ion tolerance using fully tryptic specificity (KR|P) permitting up to two missed cleavages. Search results were filtered to a 0.6% PSM-level (Peptide to Spectrum Match-level) FDR using Percolator (version 3.1), which we determined in this experiment to closely track to a 1% peptide-level FDR. Site localization was performed using an in-house implementation of Ascore that was modified to not compete localization forms against each other in order to have a higher chance of detecting overlapping positional isomers. We set Ascore to use a 0.02 Da fragment ion tolerance and we filtered for phosphopeptides with at least one corresponding PSM that produced an Ascore value >= 20 (p-value<0.01).

### DIA / PRM Acquisition and Processing

The Thermo Q-Exactive HF was configured to acquire 20 MS/MS scans at 30,000 resolution, AGC target 1e6, maximum inject time 55 ms, using overlapping precursor isolation windows of 20 m/z units and centered at: [500.4774, 520.4865, 540.4956, 560.5047, 580.5138, 600.5229, 620.5319, 640.541, 660.5501, 680.5592, 700.5683, 720.5774, 740.5865, 760.5956, 780.6047, 800.6138, 820.6229, 840.632, 860.6411, 880.6502, 900.6593, 490.4728, 510.4819, 530.491, 550.5001, 570.5092, 590.5183, 610.5274, 630.5365, 650.5456, 670.5547, 690.5638, 710.5729, 730.582, 750.5911, 770.6002, 790.6093, 810.6183, 830.6274, 850.6365, 870.6456, and 890.6547]. Full MS scans (mass range 485-925, resolution 30,000, AGC target 3e6, maximum inject time 100 ms) were interspersed every 18 scans. MS/MS scans were programed with normalized collision energy of 27 and an assumed charge state of +2.

For PRMs, the Thermo Fusion was configured to collect MS/MS scans corresponding to 62 precursor targets in the PI3K/AKT signaling pathway scheduled with 10-minute retention time windows using Phosphopedia. The large 10-minute window enables both scheduling from Phosphopedia without additional calibration runs and the detection of alternate positional isomers that may elute far away from the target. Full MS scans (mass range 400-1600) were collected at 60,000 resolution to hit an AGC target of 3e6. The maximum inject time was set to 100 ms. MS/MS scans were collected at mass range of 100-1600, resolution of 30,000, AGC target of 1e6, and maximum inject time of 55 ms. The isolation width was set to 0.7 m/z with a normalized collision energy of 27.

A Bibliospec (1) HCD spectrum library of tryptic phosphopeptides was created from the Thermo Q-Exactive data previously published in Lawrence *et al.* (2) using Skyline (version 3.1.0.7382) (3). This .BLIB library and accompanying .iRTDB normalized retention time database were used to search the .mzMLs for peptides. Thermo .RAW files were converted to .mzML and .mzXML formats using the ProteoWizard package (version 3.0.10922) where they were peak picked using vendor libraries and deconvoluted using Prism in “overlap_only” mode. We used an in-house library search engine to detect peptides in a peptide-centric approach. Our engine searches DIA data using b- and y-ion fragments of charges +1 and +2, and includes phosphate neutral losses that can be found in library spectra. We applied the following settings: 30,000 resolution (effectively +/− 16.7 ppm tolerance) for precursor, fragment, and library. Detected features were assigned and corrected to <0.01 FDR using Percolator version 3.1. We also used DIA-Umpire to extract peptide signatures using the same DIA files after overlap-deconvolution.

DIA-Umpire was configured to use 10 ppm mass tolerances and to extract +2 to +4 charged peptides. DIA-Umpire produces three .MGFs for each .mzXML; all three were searched separately with Comet and the results were combined together for Percolator and Ascore interpretation. All Comet, Percolator and Ascore settings were identical to those used for the DDA experiments.

### Thesaurus Scoring

Site-specific fragment ions can distinguish phosphoisomers but applying current site localizing tools originally designed for DDA (4) to DIA can be problematic because they assume a constant background noise level inconsistent with the high level of background interference ions found in DIA data (Supplementary Figure 1). We account for this by calculating a background frequency distribution to estimate the likelihood of detecting an interfering signal as the frequency of each m/z in the raw file for a given precursor isolation window. Our approach allows us to quickly query the distribution of ions for every fragment ion individually, enabling the use tight mass tolerances (measured in ppm) to assess the likelihood of interference.

For each queried phosphopeptide, we determine the set of combinatorial permutations corresponding to the number of phosphorylations and the number of potential phosphoacceptor sites. Starting from the two outside positional forms (where the phosphates are closest to the peptide N- or C-terminus), we extract chromatograms for fragment ions in a window +/− 10% of the total acquired chromatographic time from the retention time anchor. At every retention time point in that window we calculate a score (Thesaurus score) based on the X!Tandem HyperScore where the function is the dot product of the intensities in the acquired spectrum (*I*) and the library spectrum (*P*) multiplied by the factorial of the number of matching ions:

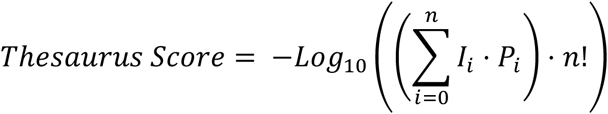

If no library spectrum is available (i.e. that phosphopeptide isoform permutation is not present in the library) then a synthetic library spectrum is generated from the anchor by shifting fragment ion peak intensities for each B-type, Y-type, +2H, and neutral loss ions to the appropriate M/Zs, using an approach similar to that used in SpectraST (5). The score vector is smoothed across time using a moving average weighted by a Gaussian based on a parameterized average retention time peak width. At the maximum scoring point we calculate a p-value as the probability of finding all of the detected site-specific ions (*n*) by chance from the background frequency distribution (*m*) and the total number of site-specific ions considered (*N*). The localization score is the -log_10_(p-value) to produce a positive score for higher confident localizations:

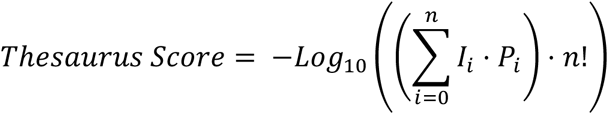

This score is also smoothed across time by Gaussian weighting.

Thesaurus then extracts the chromatographic shape of the localizing fragment ions to calculate the total number of co-eluting fragments, defined as those fragments match the peak shape with Pearson’s correlation coefficients greater than 0.75. A phosphoisomer is considered localized if the apex p-value is below a user settable threshold (typically p <=0.01 or localization score >=2) and at least three fragment ions match the shape derived from only the localizing fragment ions.

If we can localize the phosphoisomer in question, we proceed to find other internal localized forms. Because these forms share site-localizing fragment ions, Thesaurus cannot assign internal forms to the same retention time as previously detected forms. Thesaurus uses a fragment ion “exclusion list” to prevent the use of previously considered site localizing ions within the retention time window of that peak.

Finally, if we cannot assign a retention time to the sites in question with the p-value threshold, we iteratively search for an increasingly ambiguous localization that can provide some site localization-range information. For example, when localizing singly phosphorylated SGSVSNQR, we consider:

> localized: (S[+80])GSVSNQR and SGSV(S[+80])NQR
>
> ambiguous: (SGS[+80])VSNQR and SG(S[+80]VS)NQR
>
> Similarly, for doubly phosphorylated SGSVSNQR, we consider:
>
> localized: (S[+80])G(S[+80])VSNQR and SG(S[+80])V(S[+80])NQR
>
> ambiguous: (S[+80])G(SVS[+80])NQR, (SGS[+80])V(S[+80])NQR,
>
> (S[+80])G(S[+80]VS)NQR, and (S[+80]GS)V(S[+80])NQR

A parallel process is performed using decoy-generated spectra and the scoring features for both are fed into Percolator 3.1 to generate Q-values. Localizations are additionally filtered to a user-settable Q-value (typically 0.01) to ensure high confidence detections.

Positional isomer searching can be performed in an “uncalibrated” manner (where retention times in the library are assumed to be precise) if the DIA data was searched directly with a DIA library search tool or if DDA experiments were run concurrently (SWATH). Alternatively, Thesaurus supports searching in a “calibrated” manner, which assigns relative retention time ordering by searching each peptide anchor across the entire experiment window using the Thesaurus score if retention times are unknown (e.g. importing NIST libraries) or need to be calibrated (e.g. with spectrum libraries acquired on different platforms, gradients, or HPLC columns). Alternatively, searches can be performed across the entire acquisition window when analyzing targeted PRM data. We have also enabled options for only calculating localization scores and estimating FDR, skipping the detection of positional isomers that are not found in the library.

### Quantitation and Statistical Analysis

We used strict criteria to consider a localized peptide quantifiable. In addition to the localization scoring requirements, we also required at least three quantitative fragment ions and that the localized form was observed in every replicate of at least one condition. Thesaurus uses the site specific fragment ions to determine the shape of the peak and assigns quantitative fragment ions as those that match that shape for quantification with Pearson’s correlation coefficients greater than 0.9. Quantification was performed by summing the background-subtracted peak areas of site-specific fragment ions or all fragment ions, depending on the level of peptide separation. Background subtraction removes the trapezoidal area below the peak integration window. Integrated intensities were normalized within each replicate group, and across groups to the control intensity median. Statistical analysis was performed via 2-tailed t-tests paired within each replicate for individual sites, or globally with Benjamini-Hochberg FDR corrected one-way ANOVAs.

PRM results were further validated with follow-up analysis in Skyline-Daily (version 3.6.1.10615). Skyline was configured to extract all +1 and +2 b- and y-ions, including neutral losses of phosphate, as well as precursor traces for the monoisotopic, first and second isotopes. After initially importing the runs, peptides were hand-curated to match the retention time boundaries determined by site-localizing analysis. Fragment ions that appeared to be interfered with were removed from the analysis.

### Software and Data Availability

Thesaurus is an open-source, cross-platform Java application available at http://bitbucket.org/searleb/thesaurus/. While Thesaurus can read and produce reports to enable Skyline analysis, it is a stand-alone tool and does not require any other software to run. Thesaurus is fully multithreaded and designed to work on typical desktop computers where searches of individual mzML files in this experiment took on average of 10-12 minutes to complete with commodity PC hardware. All mass spectrometry files (Supplementary Table 2) presented here have been deposited to the Chorus Project (https://chorusproject.org/) with the project identifier 1374.

